# Learning Environment-Associated Marker Rankings with Attention-Based Graph Neural Networks

**DOI:** 10.64898/2026.09.21.753280

**Authors:** Amir Morshedian, Otilia Pasculescu, Mike Domaratzki

## Abstract

Genotype-by-environment interactions can complicate plant breeding because genetic effects may vary across weather regimes, locations, and years. Although genomic prediction models can use dense marker data to predict complex crop traits, they often provide limited information about how marker relevance changes across environmental conditions. This study introduces a graph-based framework for estimating environment-associated marker rankings from multi-environment crop trial data. We introduce an attention-based graph neural network that jointly represents SNP markers and weather-based environment vectors as interacting nodes and is designed to learn environment-associated marker rankings from supervised yield data. The model is trained using yield prediction as the supervised objective, and the learned marker-level attention signals are aggregated to estimate marker rankings overall and within groups of similar environments. We evaluate the framework on two genotype-by-environment datasets. In both datasets, we examine whether learned marker rankings are stable across independent training runs and determine the overlap between high-ranked markers and external GWAS references. In a maize dataset with data from 212 different site-years, we rank markers separately within each weather-defined environment cluster and define a global marker set as the markers shared across all clusters. This provides a structured way to examine how marker rankings vary across weather-defined groups and which markers remain important across environments. We further encode each weather variable with a separate LSTM combined by attention, which makes the contribution of individual weather variables readable, and compare the resulting variable ranking against an independent SHAP-based attribution. Genes located near the top-ranked markers are also examined for functional enrichment. Overall, the results suggest that graph-based, environment-conditioned marker attribution may complement genomic prediction as an exploratory tool for studying context-dependent genomic signals in multi-environment trials.

## 1 Introduction

Crop performance depends on both genetic background and growing environment. In multienvironment trials, the same genotype can perform differently across location, and weather conditions. This variation is important for breeding because farmers need cultivars that perform well under realistic and changing environments. Climate variability explains a substantial part of yield variation in major crops, and breeding programs increasingly use genomic and environmental data to identify promising genotypes [1, 2, 3].

Genomic prediction uses genome-wide marker information to predict complex traits and has become an important tool in plant breeding [4, 5, 6]. However, genotype-by-environment interaction remains a central difficulty. Prior work has shown that marker effects and crop performance can differ across environmental conditions, which limits the usefulness of models that treat marker relevance as fixed across all environments [7, 8].

Several genomic prediction models have been developed to include environmental information. Reaction-norm models and marker-by-environment interaction models use environmental variables, crop-stage summaries, or weather-derived kernels to improve prediction across environments [9, 10, 11]. More recent work has used large multi-environment crops datasets, including the Genomes to Fields project, to study yield prediction with genotype, weather, soil, and management information [12, 13, 14].

Most of this work is evaluated mainly through prediction accuracy. This is critical but it does not directly answer which markers are most relevant under different environmental contexts. Marker discovery is often studied with GWAS or QTL analysis [15], while genomic prediction models are often treated primarily as prediction tools [5, 6].

In this work, we study environment-associated marker ranking using a novel graph neural network with attention. The model represents SNP markers and weather-based environment vectors as interacting nodes. Yield prediction is used as the supervised training objective. After training, we analyze marker rankings globally, across independent training runs, and, when sufficient environments are available, within groups of similar weather conditions. We treat these rankings as model-based attribution scores and evaluate them using reproducibility and external biological references.

We study four questions: (i) can a supervised machine learning model learn useful marker rankings from genotype and weather data; ii) do marker rankings change across groups of similar weather conditions; (iii) do high-ranked markers overlap with established GWAS loci and genes with related functional annotation; and (iv) which weather variables does the environment encoder rely on, and do independent attribution methods agree on them? We evaluate these questions on two yield-focused genotype-by-environment datasets. In the dataset with greater environmental diversity, we additionally rank markers separately within weather-defined environment clusters and define a global marker set as the markers shared across clusters.

Our contributions are as follows:

- We formulate environment-conditioned marker ranking as a supervised machine learning problem in genotype-by-environment analysis and, to our knowledge, present the first crop-study application of an attention-based graph neural network designed explicitly for this task in multi-environment trials.
- We evaluate the learned rankings using reproducibility across independent runs, comparison across weather-defined environment groups, overlap with external GWAS loci, and functional annotation of nearby genes.
- We extend the environment encoder to provide interpretability at the level of individual weather variables, and cross-check the resulting variable ranking against an independent SHAP-based attribution.
- We show that graph-based, environment-conditioned marker attribution can be used as an exploratory complement to genomic prediction in multi-environment crop trials.

## 2 Related Work

### Environment data in genomic prediction

Recent studies have used environmental information in several ways for multi-environment genomic prediction. Feature-engineered environmental covariates have been used to improve genomic-enabled prediction [16], while enviromics matrices have been proposed for mixed models [17]. Nonlinear kernels with envirotyping data can improve genome-based prediction in multi-environment trials [18]. Other work has modeled G*×*E using historical weather information, envirome-wide association analysis, and weighted kernels for multi-environment prediction [19, 20, 21]. Environmental covariates have also been used outside yield prediction, including plant disease prediction [22]. Large-scale enviromic modeling has also been used for environment-associated variety recommendation [23]. ECGC introduced a statistical framework that searches for environmental covariates associated with genotype-by-environment interactions by correlating environmental similarity with genetic correlations across environments and then performing association mapping on estimated sensitivities [24]. Biclustering-based approaches have also been used for crop phenotype prediction by explicitly explaining genotype-by-environment interactions through genotype–environment blocks with approximately additive structure [25]. Most of these studies rely on non-deep-learning approaches.

### Machine learning and deep learning for genomic prediction

Machine-learning and deep-learning models have also been applied to genomic prediction with multiple data sources [26]. Machine learning has been used to combine genetic and environmental information for maize grain yield prediction [27]. Deep-learning frameworks have combined genotype-related information and weather time series through LSTM and temporal attention for crop yield prediction [28]. Deep-learning models have integrated genetic, environmental, and management data for yield prediction, and multimodal architectures have been used to fuse genomic and phenomic inputs in wheat breeding [29, 30]. Deep-learning approaches that integrate genomic and environmental data for crop genomic selection have also been reviewed [31]. Several recent architectures are especially relevant to genotype-by-environment modeling. GxENet incorporated G *×* E into fully connected neural networks for wheat yield prediction [32], and GEFormer combined gating mechanisms with linear attention for genotype-environment genomic prediction [33]. WheatGP used CNN and LSTM modules for genomic prediction [34], while MeNet introduced a mixed-effect deep neural network for multi-environment genomic prediction [35]. Other recent deep-learning models include hybrid CNN/LSTM/ResNet architectures [36], GP-Transformer for transformer-based genomic prediction [37], GViT-GP with cross-attention and genomic relationship information [38], Cropformer as an interpretable genomic prediction model [39], and CropARNet with attention and residual modules [40]. Attention models have also been extended to multi-environment genotype–phenotype settings, showing that such architectures can model epistatic and gene–environment effects while supporting cross-environment transfer [41]. Deep k-means clustering neural networks have also been used to partition maize trial locations into small ecological regions, within which GWAS and ANOVA were applied to identify SNPs significantly associated with yield and correlated with environmental factors such as temperature and precipitation. This work targets marker discovery through association testing rather than yield prediction [42]. Recent work has also examined robustness issues in deep-learning genomic prediction using obscured and ensemble models [43].

### Graph neural networks for genomic prediction

Graph-based models have also been explored for genomic prediction, although rarely applied with environmental data. HGATGS introduced a hypergraph attention network for genomic selection [44]. GCN-RS used sub-sampling graph neural networks for genomic prediction of quantitative traits, using graphs derived from genomic relationships among individuals [45]. Bipartite graph neural networks have also been proposed for maize yield prediction with trait-missing data, in which one node type represents individual field-trial records (a specific variety-location-year combination) and the other represents measured variables, including meteorological factors and agronomic traits; edges are weighted by the observed value of a feature for a sample, with missing values represented as zero-weighted edges to be imputed [46]. Other graph-based approaches have been applied to corn yield prediction under attribute-missing data and to genotypic-topological yield prediction with GraphSAGE [47, 48]. These studies show that graph-based learning can be effective for genomic prediction, but they are evaluated primarily as prediction models and do not explicitly focus on environment-conditioned marker ranking. The work is also connected to prior graph-based and attention-based genotype–environment modeling for maize yield prediction [49].

## 3 Method

The goal of this study is to learn marker-importance rankings that depend on both genotype and environmental information. For each dataset, we consider three inputs: a genotype marker matrix, daily weather records collected across the growing season for each environment, and observed yield values for genotype–environment pairs. Let *g* denote a genotype and *e* denote an environment. The pipeline consists of four main steps. First, each genotype is represented by a marker sequence

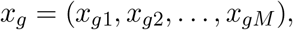

where *x*_*gm*_ is the coded value of marker *m* for genotype *g*. Second, the daily weather sequence for each environment is encoded into a fixed-length vector using an LSTM, producing an environment representation *z*_*e*_ for each environment. Third, the marker sequences and environment vectors are passed to a graph attention network (GNN), in which marker nodes and environment nodes exchange information through attention-based message passing. The model is trained using a supervised yield-prediction loss, which serves primarily as a learning signal for estimating marker importance. Finally, after training, marker-level attention weights are extracted from the model and aggregated into a global ranking. Fig. 1 summarizes the full pipeline. The following subsections describe these components in more detail.

**Fig 1:**
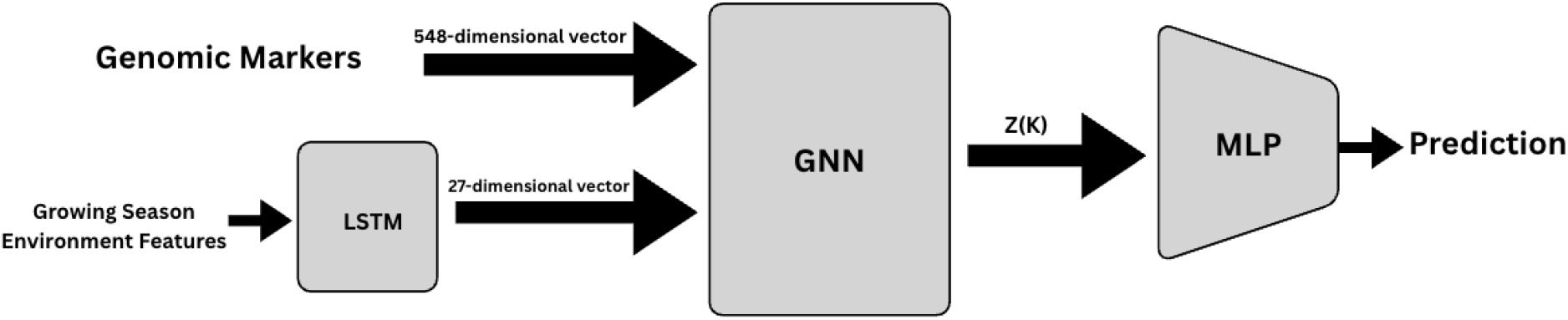
Overview of the marker-ranking pipeline. A genotype marker matrix and daily weather records across the growing season are used as inputs. Weather sequences are summarized into fixed-length environment vectors using an LSTM. The genotype marker sequences and environment vectors are then passed to a graph attention network, which is trained using supervised yield prediction. After training, marker-level attention weights are extracted and aggregated across genotype–environment observations to produce a global marker-importance ranking.

### 3.1 Weather Sequence and LSTM Environment Embedding

Each environment is represented as an ordered sequence of daily weather records collected across the growing season. For environment *e*, we write

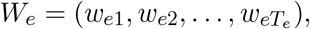

where *T*_*e*_ is the number of recorded days and *w*_*et*_ ∈ ℝ^*F*^ is the weather feature vector for day *t*. In our implementation, each daily vector contains *F* = 5 weather variables selected from an initial set of 16 candidates; details of this screening step are provided in Appendix A.

The daily weather sequence is summarized using a single-layer unidirectional LSTM. Let *h*_*et*_ and *c*_*et*_ denote the hidden state and cell state for environment *e* at day *t*. The LSTM update is

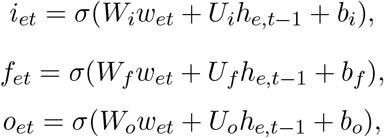

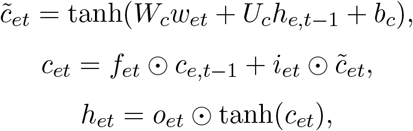

where *i*_*et*_, *f*_*et*_, and *o*_*et*_ are the input, forget, and output gates, respectively,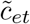 is the candidate cell state, *W*_*i*_, *W*_*f*_, *W*_*o*_, *W*_*c*_ are input-to-hidden weight matrices, *U*_*i*_, *U*_*f*_, *U*_*o*_, *U*_*c*_ are hidden-to-hidden weight matrices, *b*_*i*_, *b*_*f*_, *b*_*o*_, *b*_*c*_ are bias vectors, *σ*(·) is the logistic sigmoid and ⊙ denotes element-wise multiplication.

The final hidden state is used as the environment embedding,

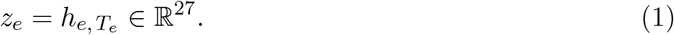

That is, for each environment *e*, we take the hidden state at its last recorded day *T*_*e*_ and use it as the fixed-length environment vector. Thus, each environment is summarized by the embedding in Eq. 1, which is used as the environment-side input to the graph attention model. Previous work [49] showed the value of tuning the environment-vector dimension; Appendix B reports the comparison used here to select dimension 27.

### 3.2 Graph Attention Network for Marker Ranking

For each genotype *g*, the GNN takes as input its marker sequence *x*_*g*_ together with the environment embeddings *z*_*e*_ from Eq. 1 for the environments in which that genotype was observed, denoted by *E*(*g*). The genotype input is kept at the marker level rather than reduced to a low-dimensional summary, so that each marker remains available for downstream ranking. The model represents markers and observed environments as nodes and applies attention-based message passing between them. Thus, each genotype-specific input contains all *M* markers, but only the environments in *E*(*g*). Each marker node is initialized from a marker-specific representation that combines its coded SNP value with its marker identity.

The marker values and environment vectors are first projected into a common hidden space. At layer *ℓ*, each observed environment node attends over the marker nodes. The environment-to-marker attention score is

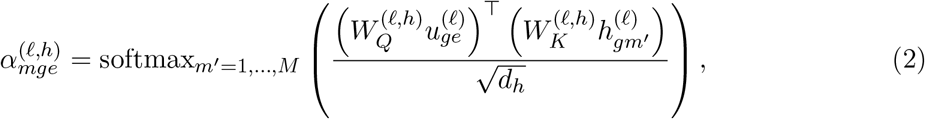

where 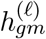 and 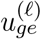 are the marker and environment node states at layer *ℓ, h* is the attention-head index, and *d*_*h*_ is the head dimension. Eq. 2 defines the environment-to-marker attention mechanism. These attention operations update both marker-node and environment-node states. After *L* layers, the model produces updated marker states 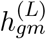 and environment states 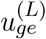.

For each observed genotype–environment pair (*g, e*), the final environment-to-marker attention is averaged across heads,

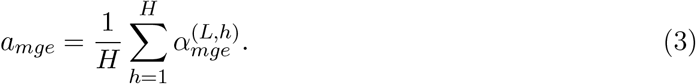

Here, the final attention score in Eq. 3 measures how much marker *m* is attended to for genotype *g* in environment *e*. The model uses these marker-attention scores to form the environment-associated genotype summary,

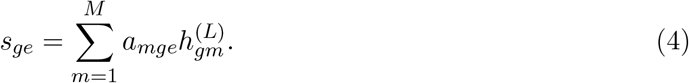

Eq. 4 defines the environment-associated genotype summary. This summary gives larger contribution to marker states with higher attention. It is concatenated with the final environment-node state and passed to the feed-forward prediction head:

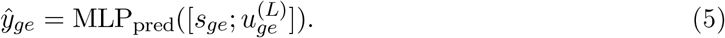

Eq. 5 defines the feed-forward prediction head. Let *q*_*ge*_ denote the normalized replicate-count weight for genotype–environment pair (*g, e*). The model is trained on observed pairs using the weighted loss, which combines mean squared error and mean absolute error,

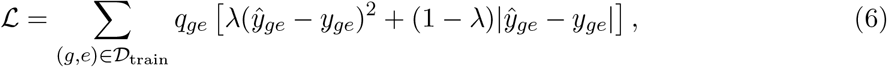

where *y*_*ge*_ is the observed yield and *λ* controls the relative weight of the two loss terms. Eq. 6 defines the training objective.

After training, the learned GNN is applied again to all observed genotype–environment pairs to extract one marker-attention vector

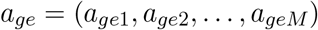

for each pair. These vectors are then aggregated across observations to obtain the global importance score for each marker,

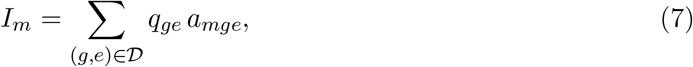

where *a*_*mge*_ is the attention weight assigned to marker *m* for genotype *g* in environment *e*, *D* is the set of observed genotype–environment pairs, and *q*_*ge*_ is a normalized observation weight. Eq. 7 defines the global marker importance score. This aggregation provides an overall summary of marker relevance across observed genotype–environment pairs, while environment-conditioned differences are examined separately in the cluster-specific analyses. In our implementation, *q*_*ge*_ is obtained by normalizing the replicate count for the pair (*g, e*), so pairs with more replicates contribute more to the final marker score. Markers are then ranked in decreasing order of *I*_*m*_, and the top-ranked sets are used in the downstream analyses.

## 4 Experiments

### 4.1 Datasets and Global Marker Ranking

We evaluate the proposed marker-ranking framework on two yield-focused genotype–environment datasets. The maize dataset [50] uses a targeted 1000-genotype analysis set selected from a larger 4,928-genotype panel by taking the 800 genotypes with the highest mean observed yield among valid records and adding 200 randomly sampled genotypes from the remaining valid pool. This top-800 plus random-200 policy defines a computationally feasible, yield-enriched analysis set while retaining additional genotype diversity beyond the highest-yield group. For SoyNAM [51], we use a six-environment version after filling two missing weather environments from NASA POWER [52] and use all valid hybrids with genotype, phenotype, and weather support. Summary statistics for both datasets are reported in Table 1. Additional details on genotype input construction, including the maize marker-reduction procedure from 437,214 source markers to the final 4,050-marker matrix, are provided in Appendix A.

**Table 1:**
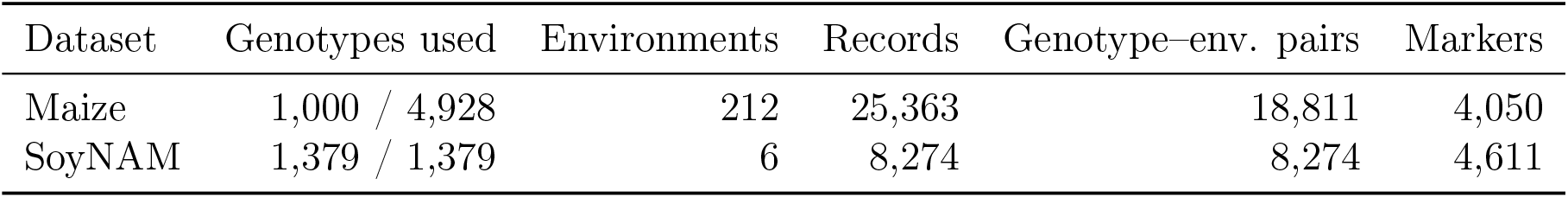
Datasets used for global marker-ranking experiments.

| Dataset | Genotypes used | Environments | Records | Genotype-env. pairs | Markers |
| --- | --- | --- | --- | --- | --- |
| Maize | 1,000 / 4,928 | 212 | 25,363 | 18,811 | 4,050 |
| SoyNAM | 1,379 / 1,379 | 6 | 8,274 | 8,274 | 4,611 |

For each dataset, daily weather records across the growing season are summarized into fixed-length environment vectors, and the proposed GNN is trained using observed yield as the supervised signal. After training, we extract marker-level attention scores and aggregate them into one global score for each marker. Markers are then sorted by decreasing score to produce a global ranking. In the downstream analyses, we focus on the top 200, top 500, and top 1,000 markers.

Prediction metrics are reported only as a training check, since the main goal here is marker ranking rather than yield benchmarking. In the maize dataset, the global model reached validation *R*^2^ = 0.61. In the SoyNAM dataset, the global model reached validation *R*^2^ = 0.66. The corresponding training summary is reported in Table 2.

**Table 2:** Training summary for the global marker-ranking models.

| Dataset | Validation MSE | Validation MAE | Validation $R^2$ |
| --- | --- | --- | --- |
| Maize | 2.94 | 1.32 | 0.61 |
| SoyNAM | 0.40 | 0.49 | 0.66 |

Fig. 2 shows the global marker-weight curves for the two datasets, with markers sorted by decreasing global marker score *I*_*m*_ from Eq. 7. In both cases, the weight mass is concentrated in a relatively small set of markers, which motivates the top-*k* analyses used in the remainder of the paper. These ranked lists define the top-*k* marker sets used in the reproducibility, GWAS-overlap, and weather-cluster analyses. The curves are descriptive only, and reproducibility of the resulting rankings is evaluated separately in section 4.2. All reported models use the same preprocessing and train/validation protocol; complete implementation and training details are provided in Table 3.

**Table 3:**
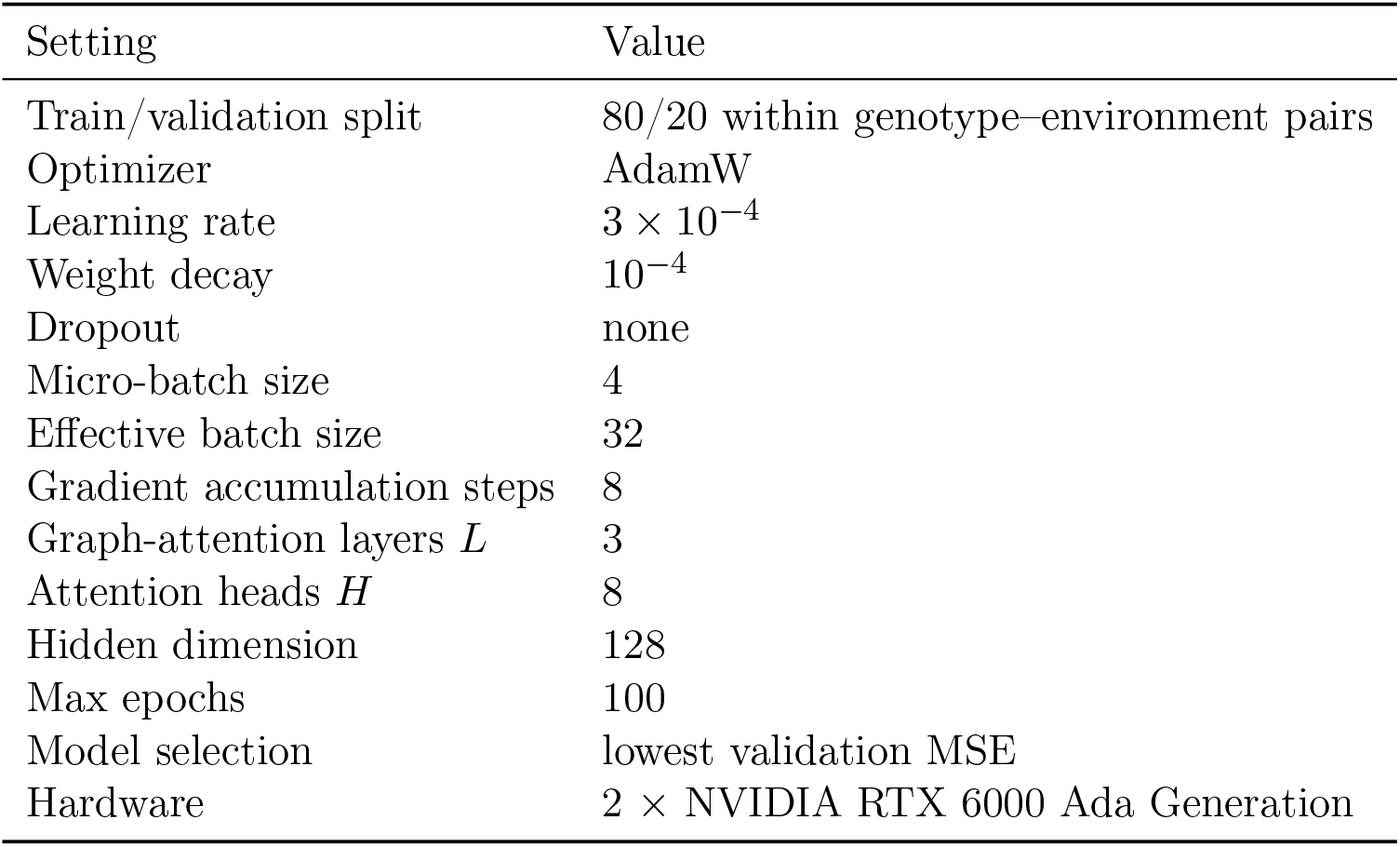
Training configuration for the GNN experiments.

| Setting | Value |
| --- | --- |
| Train/validation split | 80/20 within genotype-environment pairs |
| Optimizer | AdamW |
| Learning rate | $3 \times 10^{-4}$ |
| Weight decay | $10^{-4}$ |
| Dropout | none |
| Micro-batch size | 4 |
| Effective batch size | 32 |
| Gradient accumulation steps | 8 |
| Graph-attention layers $L$ | 3 |
| Attention heads $H$ | 8 |
| Hidden dimension | 128 |
| Max epochs | 100 |
| Model selection | lowest validation MSE |
| Hardware | $2 \times$ NVIDIA RTX 6000 Ada Generation |

**Fig 2:**
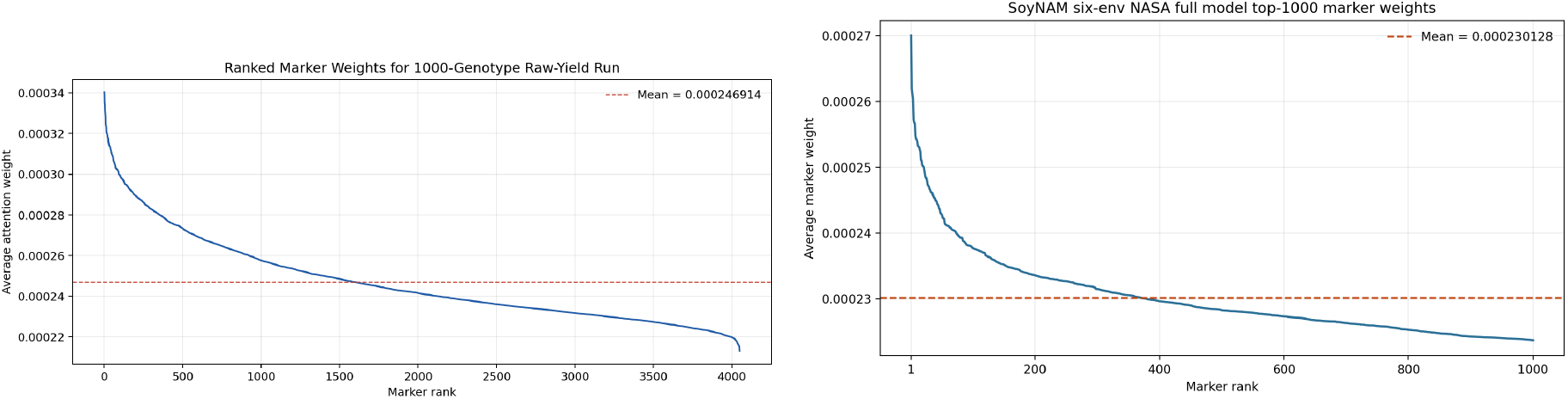
Global marker-weight curves for the maize and SoyNAM datasets. Markers are ordered by decreasing global importance score. The curves are used to motivate the top-*k* ranking analyses.

### 4.2 Reproducibility Across Independent Runs

We examine whether highly ranked markers are reproduced across repeated training runs. All 5 runs used the same phenotype data, genotype file, environment vectors, and train/validation split. For a compact reproducibility summary, we used 5 runs for each dataset and defined a marker as recurrent if it appeared in at least 3 of those 5 runs. Table 4 reports the number of recurrent markers for the top 200, top 500, and top 1000 marker sets. To calibrate these counts, we also compare them with a random baseline obtained from randomly sampled marker sets of size *k*, drawn independently across 5 runs. In both datasets, recurrence increases as the size of the reported top-*k* set increases.

**Table 4:**
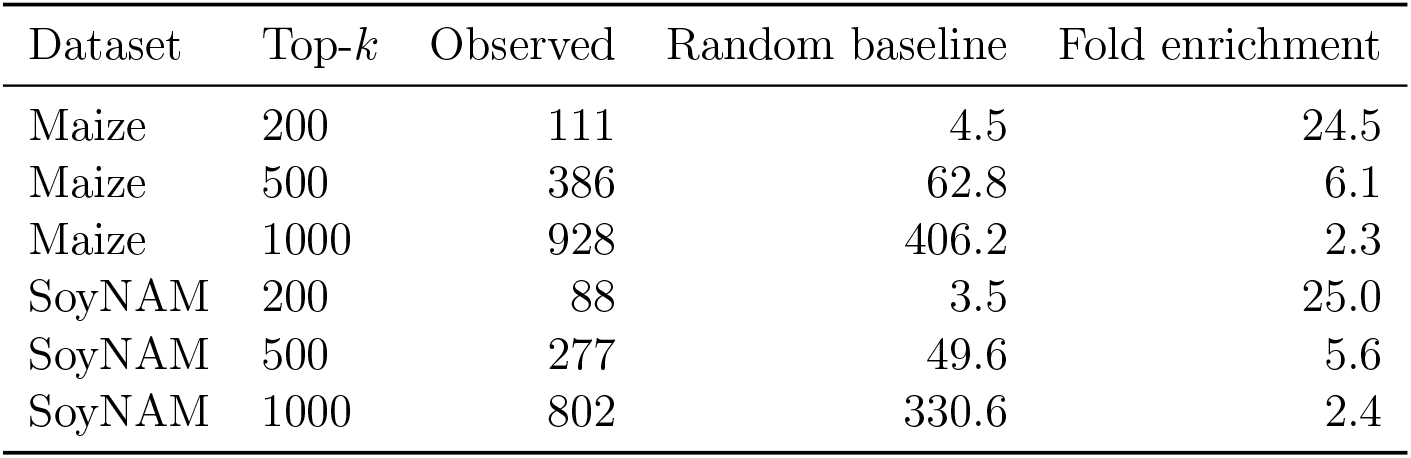
Markers shared by at least 3 of 5 independent runs. The random baseline is the expected number of recurrent markers under independent random top-*k* marker sets of the same size. Fold enrichment is the ratio of observed to expected recurrence.

### 4.3 Predictive Performance of Ranked Marker Subsets

Because the marker rankings are derived from attention weights, we evaluated whether higher-ranked markers support better prediction than equally sized random or low-ranked subsets.

Using the global maize ranking learned on the training split, we constructed top-*k*, random-*k*, and bottom-*k* marker subsets for *k* ∈ {200, 500, 1000}. For each subset, we trained a random-forest regressor with marker-only inputs to predict environment-residual yield, using the same train/validation split as in the main maize experiment. We report this analysis for maize because its larger multi-environment setting provides a more informative test set for residual-yield prediction.

Table 5 summarizes the results. Across all three subset sizes, the top-ranked marker subsets outperformed both random and bottom-ranked subsets in both weighted *R*^2^ and weighted Pearson correlation. These results shows that markers receiving higher attention scores tend to retain more downstream predictive utility than low-ranked markers and random subsets.

**Table 5:** Random-forest prediction results using marker-only inputs and environment-residual yield as the prediction target in maize. Marker subsets were defined from the GNN ranking as top-*k*, bottom-*k*, or random-*k*.

| Top-$k$ | Top $R_w^2$ | Random $R_w^2$ | Bottom $R_w^2$ | Top $PCC_w$ | Random $PCC_w$ | Bottom $PCC_w$ |
| --- | --- | --- | --- | --- | --- | --- |
| 200 | 0.01498 | 0.01314 | 0.00413 | 0.16122 | 0.14208 | 0.13976 |
| 500 | 0.01470 | 0.01324 | 0.00818 | 0.16273 | 0.15463 | 0.14849 |
| 1000 | 0.01796 | 0.01522 | 0.01155 | 0.16889 | 0.16084 | 0.15449 |

### 4.4 External GWAS Overlap

Because there is no standard benchmark that directly evaluates environment-conditioned marker rankings in this setting, we use published Genome-wide association studies (GWAS) loci as an external reference. GWAS test statistical associations between genetic markers and trait variation in a population, and they report SNPs or loci associated with the trait of interest [53, 54]. In our case, these published GWAS results provide a limited but useful reference set for checking whether the model’s top-ranked markers recover previously reported yield-related regions.

This comparison is not intended as a direct benchmark against another marker-ranking method. Instead, it serves as an external consistency check. For maize, we used the yield-related SNP sets reported by Zeng [55], Tolley [56], and Ma and Cao [57]. For soybean, we used the yield marker-trait associations reported by Diers [58]. A model marker was counted as a match if it was located on the same chromosome and within a fixed genomic window around a reference GWAS SNP. We use window-based matching because the exact GWAS SNP may not be present in our marker panel, while a nearby marker may be in linkage with the GWAS SNP. We evaluate overlap using 100 kb, 500 kb, and 1 Mb windows, with 500 kb used as the primary comparison.

As non-GNN ranking baselines, we also ranked markers by the absolute value of their Pearson correlation with observed yield and by random-forest feature importance, a widely used approach for identifying informative markers in genome-wide association data [59, 60]. For the random-forest baseline, we trained a random-forest regressor and extracted feature-importance scores for the marker features, and evaluated the resulting top-1000 marker ranking under the same GWAS-overlap criteria.

Table 6 summarizes the GWAS overlap results using the primary 500 kb window together with the univariate-correlation and random-forest baselines. Overall, these results show that the learned rankings overlap with many reported yield-related regions across several GWAS studies. Under the same matching conditions, the GNN-based ranking consistently achieves higher overlap than the univariate-correlation and random-forest baselines. Since these GWAS reference sets differ from one another, repeated overlap of the GNN-ranked markers across multiple studies provides external evidence for the learned rankings. Appendix C reports the same analysis using 100 kb and 1 Mb windows.

**Table 6:** External GWAS overlap for the top 1000 global markers using a 500 kb window around each reference SNP.

| Dataset | Reference set | Ref. SNPs | GNN overlap | GNN cov. (%) | RF baseline cov. (%) | Univariate-corr. cov. (%) |
| --- | --- | --- | --- | --- | --- | --- |
| Maize | Zeng 2022 | 59 | 28 | 47.46 | 30.59 | 23.73 |
| Maize | Tolley 2023 G2F | 21 | 12 | 57.14 | 47.62 | 33.33 |
| Maize | Ma and Cao 2021 | 22 | 8 | 36.36 | 27.27 | 18.18 |
| SoyNAM | Diers 2018 | 23 | 16 | 69.57 | 54.87 | 47.83 |

### 4.5 Weather-Defined Environment Clusters and Cluster-Specific Marker Rankings

To examine whether additional marker signals appear under different weather conditions, we performed a cluster-specific analysis on the maize dataset by ranking markers independently within each weather-defined cluster. We then defined a global marker set as the markers shared across all four clusters.

Environment clustering was performed on environment-level weather summaries using *k*-means [61]. We evaluated candidate feature-set sizes of 3, 5, and 7 weather variables together with *k* ∈ {2, 3, 4, 5}, where *k* is the number of clusters, and retained the 5-feature (see Table B), *k* = 4 solution because it gave the best silhouette score [62] in this scan. Clustering was first defined on all environments with weather data, giving 151, 72, 26, and 20 environments in clusters C0–C3, respectively. Fig. 3 summarizes the final configuration through a PC1/PC2 projection and the number of environments in each cluster. Fig. 4 shows the cluster-profile heatmap used to assign descriptive labels after clustering. The heatmap values are standardized *z*-scores. These labels were used only for interpretation, not for training.

**Fig 3:**
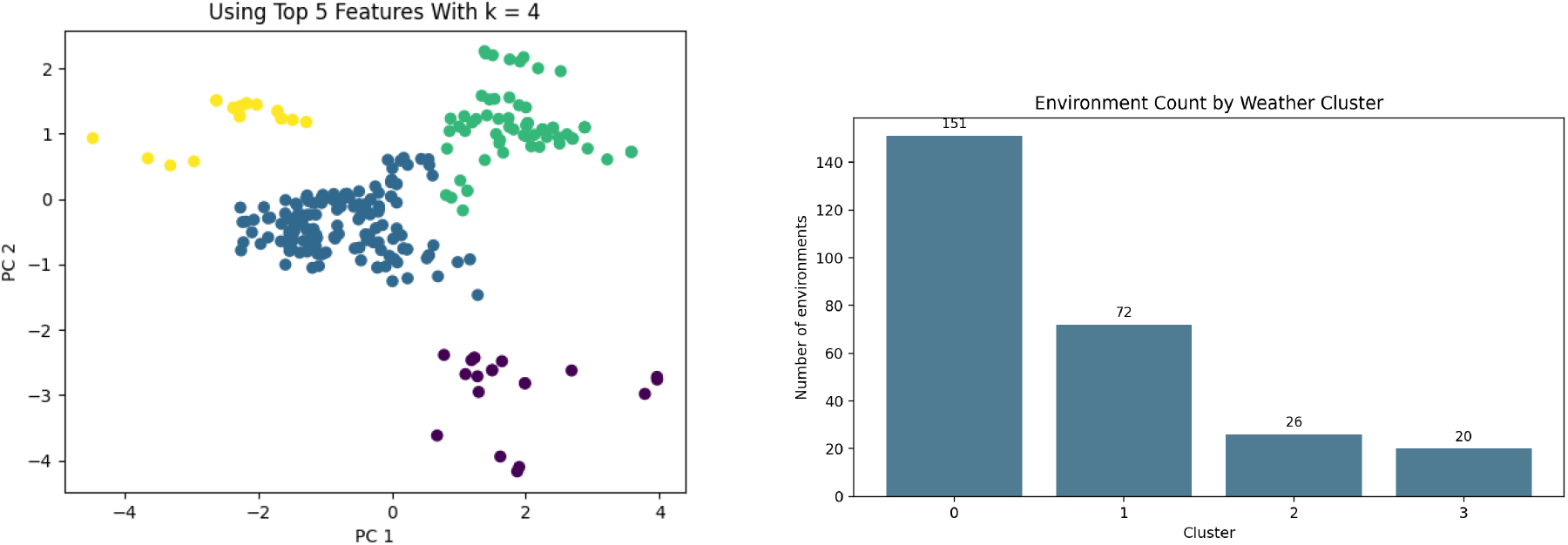
Final weather-cluster configuration for the maize environment analysis. Left: PC1/PC2 projection of the standardized five-feature environment means with the final *k* = 4 assignments. Right: number of environments in each weather cluster.

**Fig 4:**
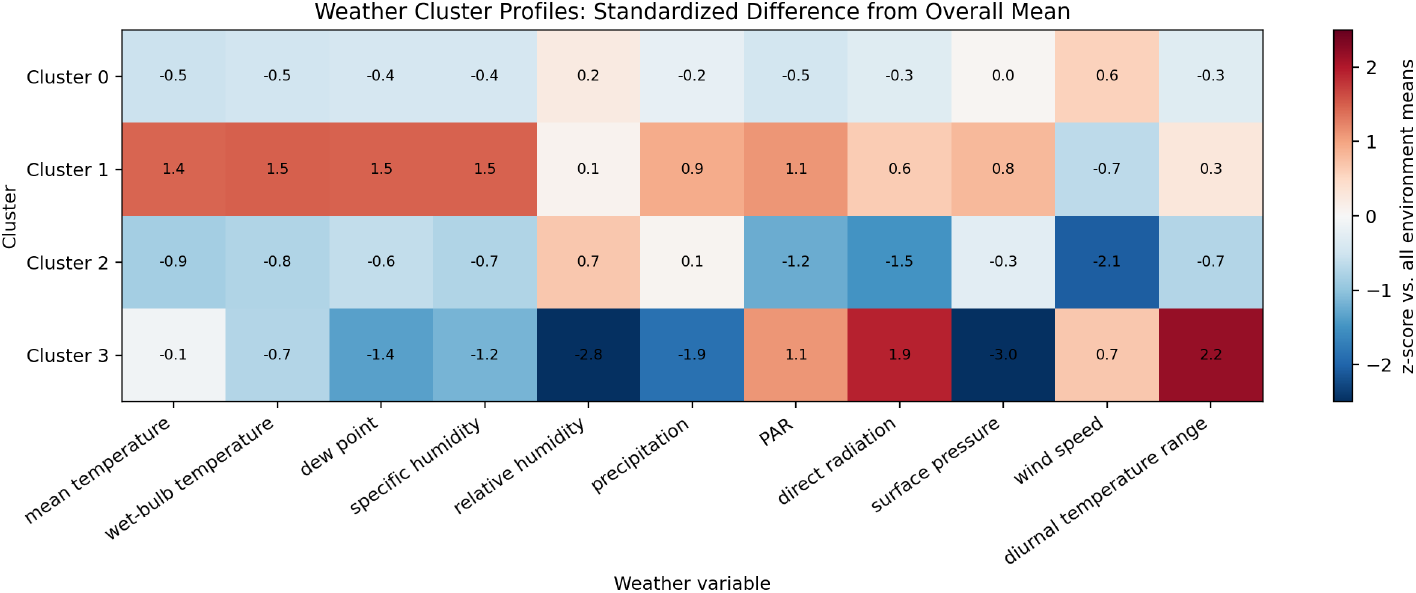
Cluster-profile heatmap for the final *k* = 4 solution. Values are shown as standardized *z*-scores relative to the full set of environments used in the clustering step. This plot was used to assign descriptive labels to the four weather clusters.

The four retained weather groups were: C0, cool-to-temperate windy conditions with moderate humidity; C1, warm humid wet conditions with low wind; C2, cool humid conditions with low radiation and very low wind; and C3, dry high-sun conditions with lower pressure and higher wind. The cluster-specific marker models were then trained only on the subset of maize environments that were available in the selected maize analysis set. After this intersection step, the cluster-specific GNN models used 18,102 (C0), 4,259 (C1), 2,930 (C2), and 4,342 (C3) phenotype records.

We then trained one cluster-specific GNN per weather group and compared the resulting marker rankings. Table 7 summarizes the overlap and uniqueness patterns.

**Table 7:**
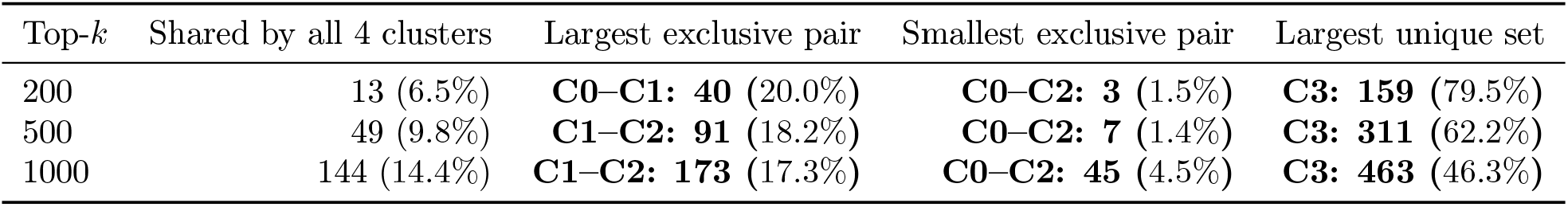
Summary of cluster-specific marker structure. All values are exclusive marker counts with the corresponding percentage of the top-*k* marker list.

Figs. 5–7 show the exact exclusive overlap structure of the cluster-specific marker sets at each top-*k* threshold. In each figure, the vertical axis lists one marker-set combination across the four weather clusters and the bar length gives the number of markers belonging exclusively to that combination, while the dot matrix on the right marks which clusters participate in each combination. Fig. 8 summarizes the pairwise overlap structure for the cluster-specific marker sets. In this heatmap, both axes index the four weather clusters and each cell reports the percentage of the top-*k* marker set shared between the corresponding pair of clusters.

**Fig 5:**
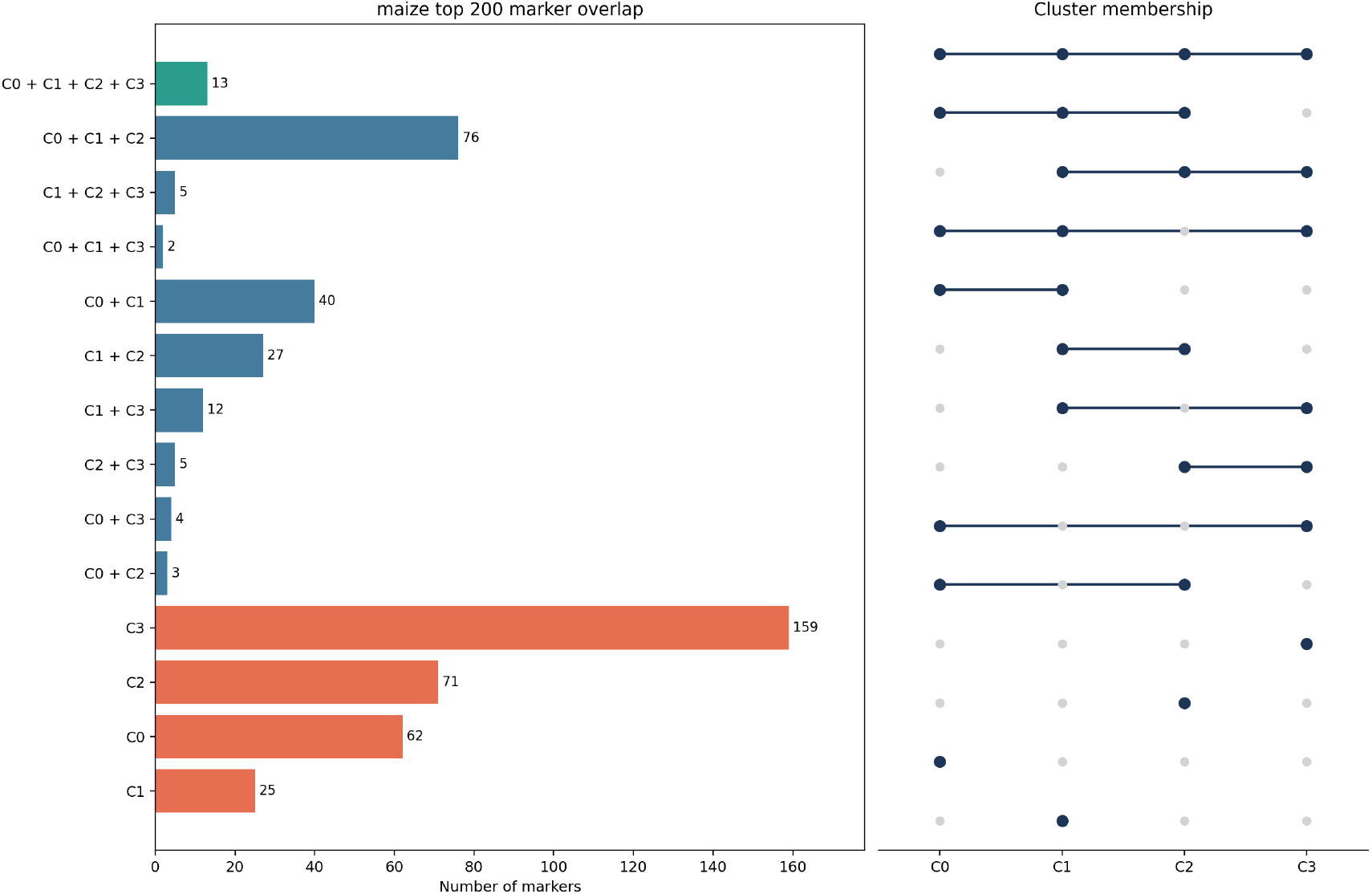
Exclusive overlap structure of the cluster-specific top-200 marker sets. Each bar gives the number of markers belonging exclusively to one combination of weather clusters, and the dot matrix on the right marks which clusters participate in each combination.

**Fig 6:**
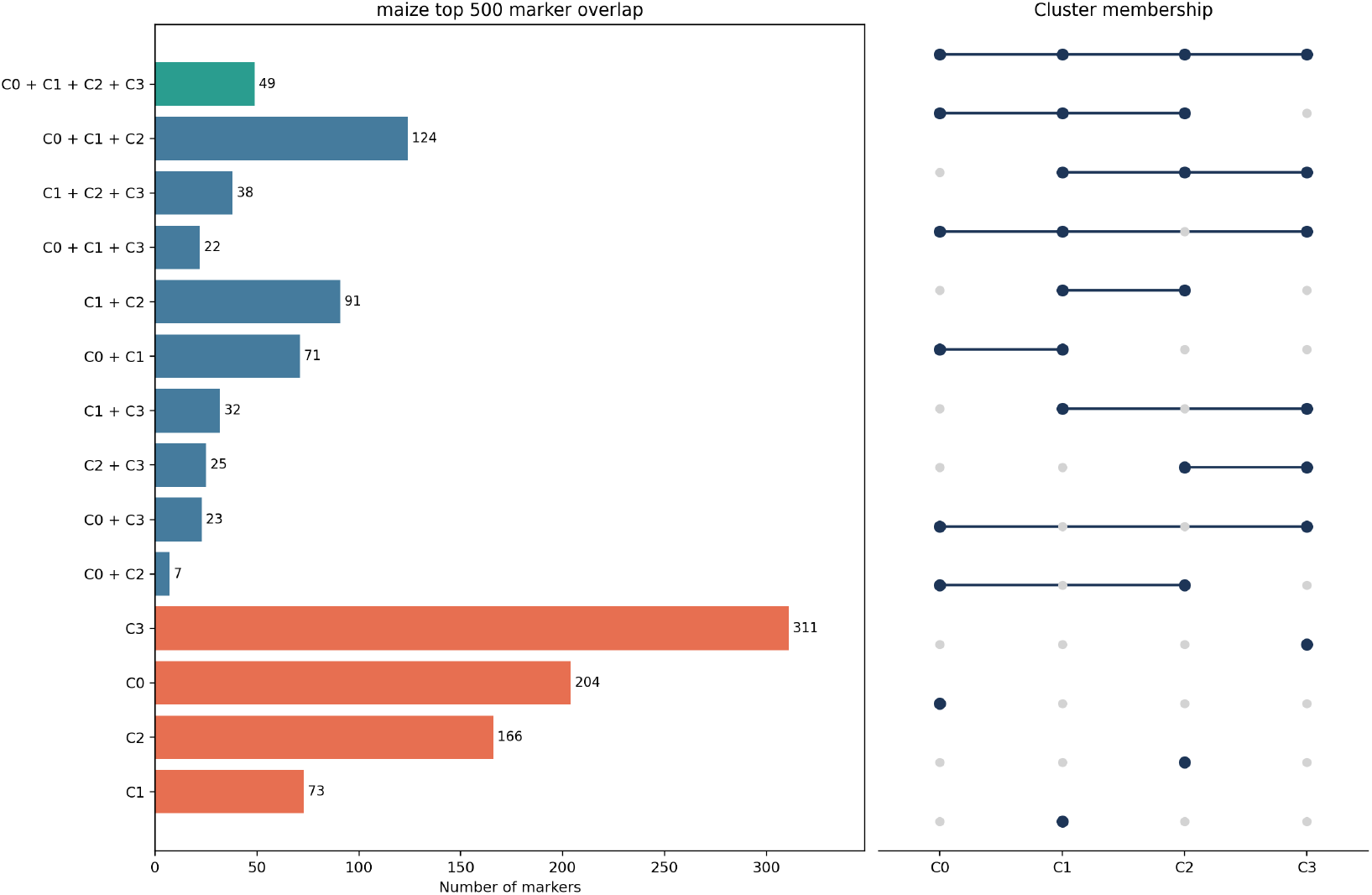
Exclusive overlap structure of the cluster-specific top-500 marker sets. Each bar gives the number of markers belonging exclusively to one combination of weather clusters, and the dot matrix on the right marks which clusters participate in each combination.

**Fig 7:**
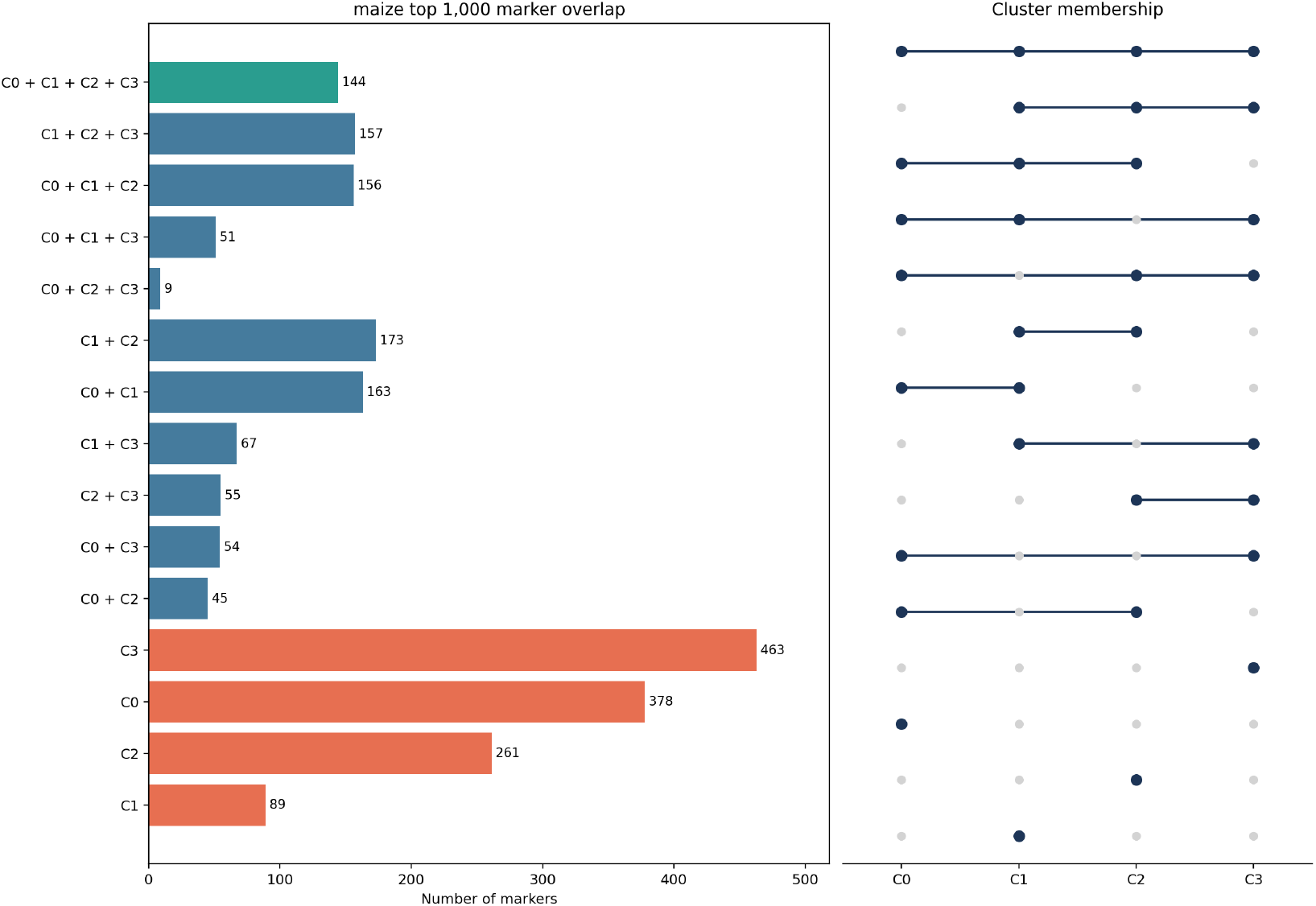
Exclusive overlap structure of the cluster-specific top-1000 marker sets. Each bar gives the number of markers belonging exclusively to one combination of weather clusters, and the dot matrix on the right marks which clusters participate in each combination.

**Fig 8:**
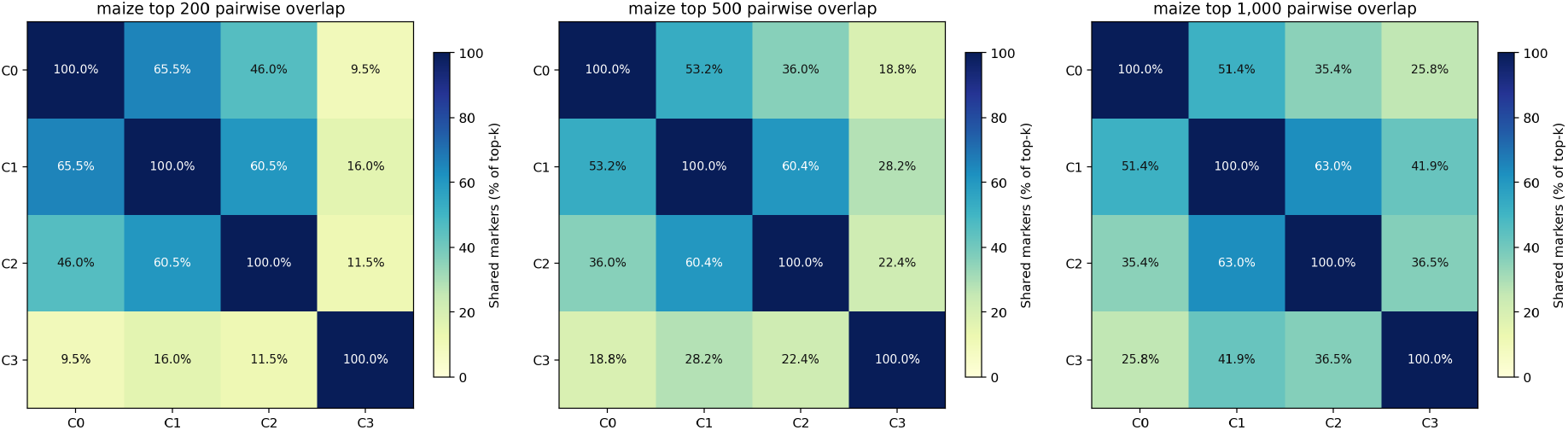
Pairwise marker overlap for the cluster-specific marker sets at top 200, top 500, and top 1000 (left to right). Rows and columns index weather clusters C0–C3, each with its own top-*k* marker set of size *k*. Each off-diagonal cell reports the percentage of one cluster’s top-*k* set that is also present in the other cluster’s top-*k* set, including markers that additionally appear in a third cluster; diagonal cells equal 100% by definition, since a cluster’s top-*k* set trivially overlaps completely with itself.

Because the cluster definitions depend on the selected weather summaries and clustering configuration, these results should be interpreted as exploratory evidence of cluster-dependent ranking differences under the current analysis pipeline.

### 4.6 Environment-Feature-Level Interpretability with Per-Variable LSTM Encoding

The environment encoder in Section 3.1 processes the five selected weather variables jointly through a single LSTM, which makes the contribution of individual variables difficult to identify. To obtain feature-level interpretability, we instead used an independent LSTM for each weather variable and combined the resulting representations through learned additive attention:

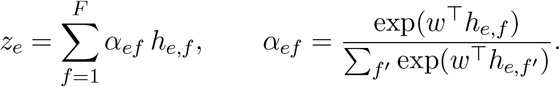

Here, *h*_*e,f*_ denotes the final hidden state of the LSTM assigned to weather variable *f* in environment *e*, and *α*_*ef*_ is the weight given to that variable in the environment representation. The attention weights sum to one across variables for each environment, so they can be read directly as the share of the representation contributed by each weather variable. All 16 candidate weather variables were used in this analysis.

To compare the attention-based ranking with an independent attribution method, the final hidden state of each per-variable LSTM was passed through a learned linear layer to obtain one scalar per weather variable. These 16 scalars were then used as input to a small MLP classifier. SHAP values [63] were computed from this classifier and used to rank the weather variables. The classifier was independently retrained 20 times, and the resulting SHAP rankings were compared with the attention ranking. Table 8 reports the stability of the SHAP ranking across these runs alongside each variable’s attention rank, and Table 9 reports the ten variables with the largest pooled SHAP attribution.

**Table 8:** Stability of the SHAP ranking across 20 independent retrainings of the attribution classifier, with each variable’s attention rank for comparison. Rows are ordered by mean SHAP rank.

| Variable | Attention rank | Mean SHAP rank | SD | Min | Max |
| --- | --- | --- | --- | --- | --- |
| RH2M | 4 | 1.50 | 0.76 | 1 | 3 |
| ALLSKY_SFC_SW_DNI | 9 | 3.15 | 2.39 | 1 | 11 |
| T2MDEW | 8 | 4.05 | 2.44 | 1 | 10 |
| T2M_MIN | 11 | 4.30 | 2.11 | 1 | 9 |
| WS2M | 2 | 4.50 | 1.40 | 2 | 7 |
| PS | 1 | 5.25 | 2.02 | 3 | 12 |
| PRECTOTCORR | 6 | 8.20 | 2.26 | 6 | 13 |
| GWETROOT | 3 | 8.30 | 1.95 | 5 | 14 |
| T2M_MAX | 15 | 9.05 | 2.09 | 4 | 12 |
| GWETPROF | 5 | 10.30 | 3.23 | 6 | 16 |
| T2M | 14 | 11.15 | 1.98 | 8 | 15 |
| GWETTOP | 12 | 11.45 | 1.96 | 9 | 15 |
| T2MWET | 13 | 12.30 | 3.06 | 3 | 16 |
| ALLSKY_SFC_PAR_TOT | 7 | 13.80 | 1.67 | 10 | 16 |
| QV2M | 10 | 14.05 | 1.64 | 10 | 16 |
| ALLSKY_SFC_SW_DWN | 16 | 14.65 | 1.53 | 12 | 16 |

**Table 9:**
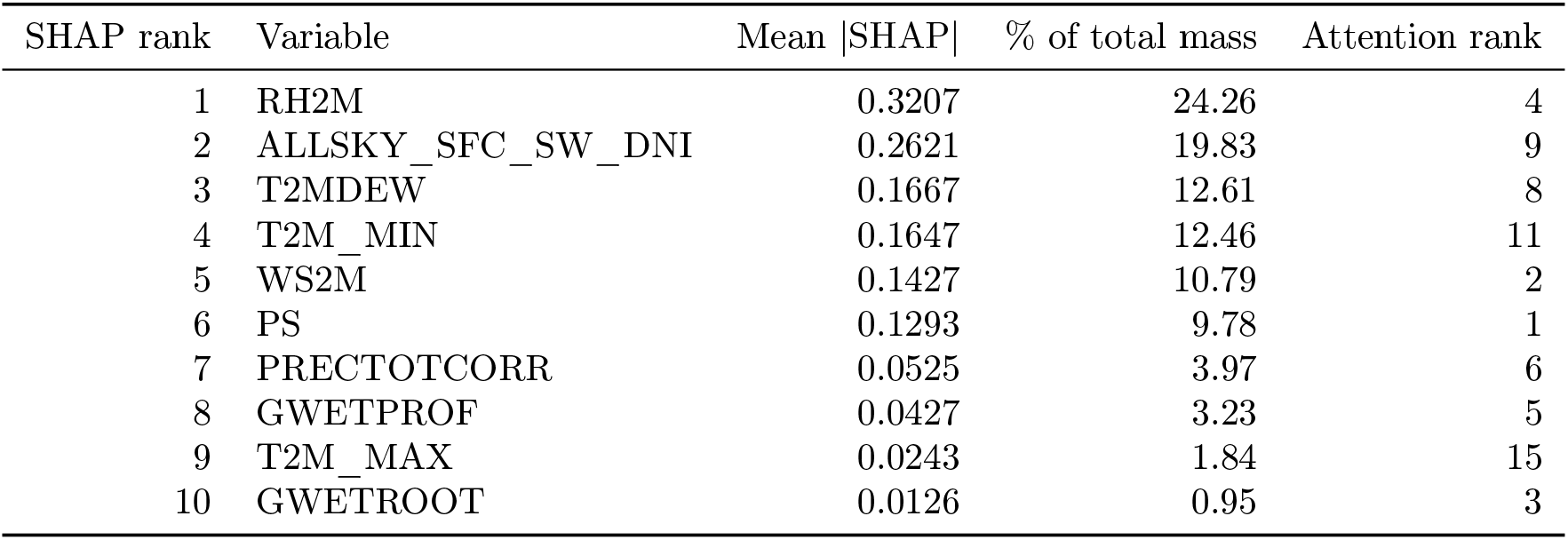
Ten highest-ranked weather variables by pooled attribution magnitude across the 20 retrainings, together with their attention rank. These ten variables account for 99.73% of the total |SHAP| mass.

The two methods identify overlapping sets of influential variables, although their orderings differ. The top-five sets overlap by an average of 2.50 variables across the 20 runs. Agreement is concentrated in a small and stable group: relative humidity (RH2M) appears in the top five of both methods in all 20 runs, followed by wind speed (WS2M) in 15 runs and surface pressure (PS) in 14 (Fig. 9). At the top-ten level, eight variables are selected by both methods (Fig. 10). Across all 16 variables the two rankings have a mean Kendall’s *τ*_*b*_ [64] of 0.356, computed with tied attributions accounted for, and the coefficient is positive in every run.

**Fig 9:**
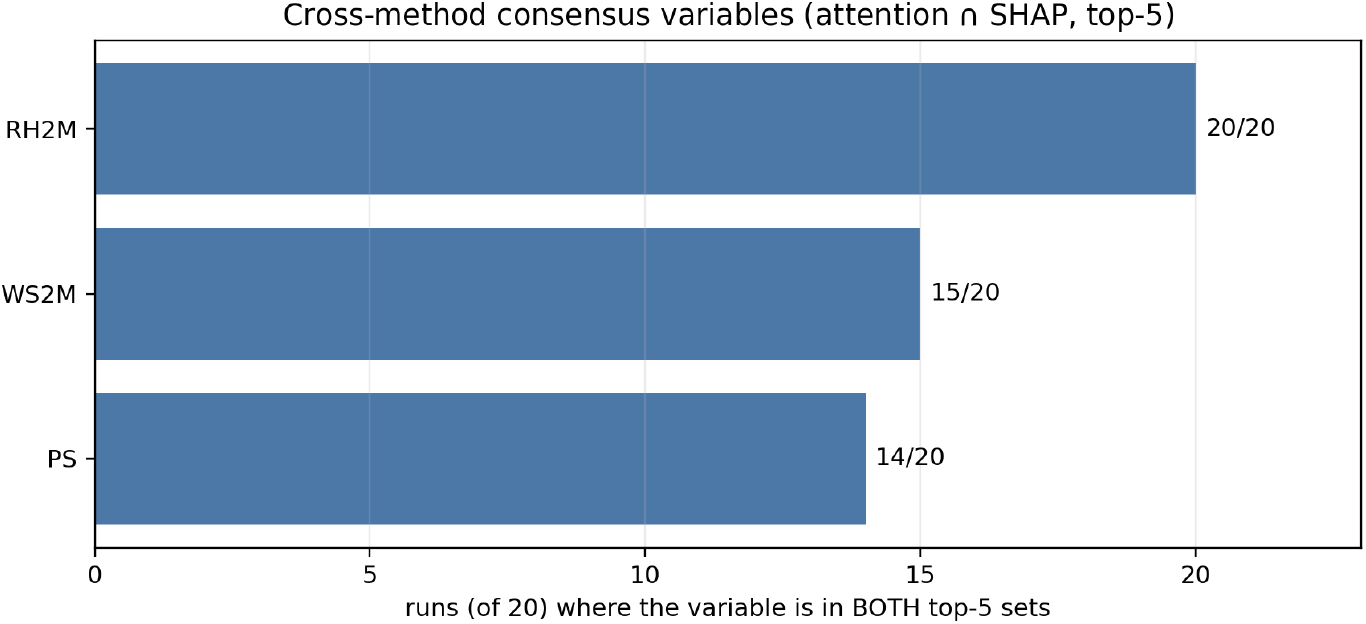
Weather variables selected by both methods. For each variable, the number of the 20 attribution retrainings in which it appears in both the attention top-five and the SHAP top-five. Relative humidity is selected by both methods in every run.

**Fig 10:**
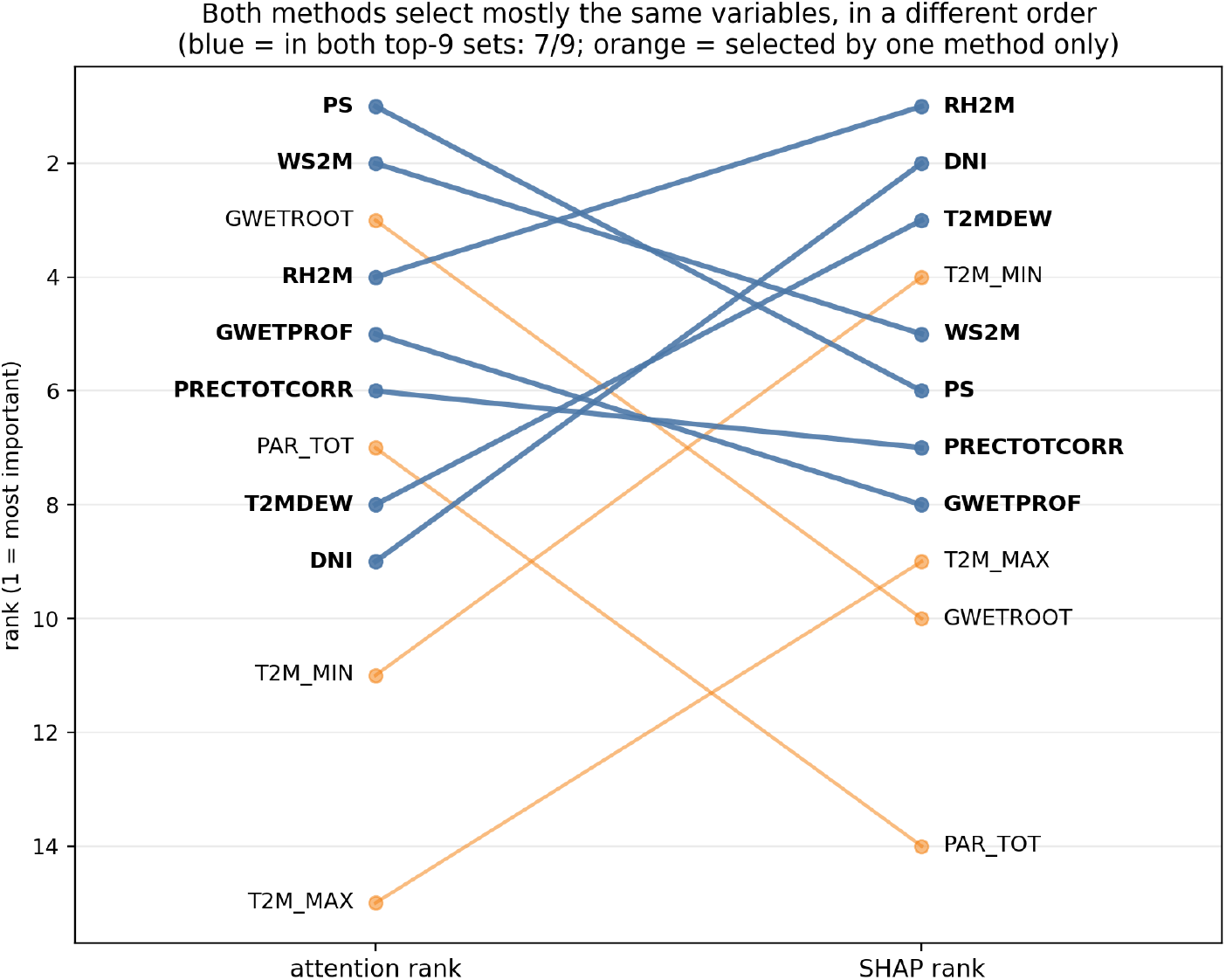
Attention rank against SHAP rank for the variables entering the ten highest-ranked positions under either method. Variables selected by both methods are shown in blue (8 of 10); variables selected by only one method are shown in orange.

The two rankings describe related but distinct quantities: the attention ranking reflects how strongly each weather variable was weighted when constructing the learned environment representation, whereas the SHAP ranking attributes the classifier’s prediction to each variable.

Agreement on which variables are influential, combined with disagreement on their precise order, is therefore the expected outcome.

### 4.7 Functional Annotation of Shared and Cluster-Specific Markers

To examine whether the ranked markers fall near genes with known biological function, each marker was mapped to the annotated genes within 500 kb using the B73 reference annotation (Zm-B73-REFERENCE-NAM-5.0) [65]. The resulting gene sets were tested for over-representation of Gene Ontology [66], InterPro [67], Pfam [68], and Plant Reactome [69] terms against the full 4,050-marker panel, using a hypergeometric test with Benjamini–Hochberg correction at a 10% false discovery rate. A single marker next to a tandem array of paralogous genes can inflate this type of analysis, so we kept only terms supported by at least three independent markers. This criterion was met by 188 of the 276 terms significant at FDR *<* 0.10 (68.1%).

For each marker set, Table 10 reports the most significant term with a known biological function. The markers shared by all four weather clusters are enriched for O-methyltransferase activity, a gene family associated with lignin biosynthesis and cell-wall composition in maize; polymorphisms in this gene family have previously been associated with biomass yield and other agronomic traits in maize [70]. The cluster-specific sets are enriched for different functional categories: an LRK10-like disease-resistance receptor kinase family in C2 (cool, humid conditions), a tryptophan-synthase-like, pyridoxal-phosphate-dependent enzyme family in C1 (warm, humid, low-wind conditions), and lipid transfer and defence-related proteins [71] in C0 (cool, windy conditions) and C3 (dry, high-sun conditions). In Arabidopsis, the corresponding tryptophan synthase *β* subunit gene coordinates plant growth and abiotic-stress responses by modulating tryptophan and abscisic-acid homeostasis [72]. The LRK10-like family identified with C2 was originally identified through its association with fungal leaf-rust resistance in wheat, and its best-characterized ortholog in Arabidopsis is instead linked to abscisic-acid signaling and drought resistance [73]; its enrichment in C2, the coolest and most humid of the four clusters, is a candidate for further investigation. Expanded results are reported in Appendix D.

**Table 10:** Functional terms enriched among genes within 500 kb of the ranked markers, restricted to terms supported by at least three independent markers (FDR *<* 0.10). Cluster-specific sets (C0–C3) use each cluster’s own top 100 ranked markers; the all-clusters set is the intersection of markers ranked in the top 1000 of every cluster. Genes is the number of genes carrying the term; Markers is the number of distinct markers located near those genes, which can be smaller than Genes when several term-carrying genes lie near the same marker.

| Marker set | Enriched term | Genes | Markers | Fold | FDR $q$ |
| --- | --- | --- | --- | --- | --- |
| All four clusters | O-methyltransferase activity | 6 | 3 | 9.9 | $3.7 \times 10^{-3}$ |
| C0 (top 100) | Lipid transfer / defence protein | 10 | 4 | 5.2 | $1.7 \times 10^{-3}$ |
| C1 (top 100) | PF00291 (Pfam domain) | 8 | 4 | 4.7 | $2.1 \times 10^{-2}$ |
| C2 (top 100) | Rust resistance kinase Lr10-like | 11 | 4 | 5.0 | $1.2 \times 10^{-3}$ |
| C3 (top 100) | Lipid transfer / defence protein | 12 | 3 | 6.2 | $2.6 \times 10^{-5}$ |

## 5 Conclusion

We introduced an attention-based graph neural network framework for learning marker rankings from genotype and weather information in multi-environment yield data. Across two datasets, the learned rankings showed reproducibility across independent runs and overlap with published GWAS regions. In maize, a random-forest evaluation showed that top-ranked marker subsets retained a modest but consistent predictive advantage over random and bottom-ranked subsets. Cluster-specific analysis further suggested that, ranking markers independently within each weather-defined cluster, the resulting rankings contain both shared and weather-associated components. Functional annotation of these marker sets showed that the shared and cluster-specific markers lie near genes enriched for different functional categories. Replacing the joint weather encoder with one LSTM per weather variable combined by attention additionally provided interpretability at the level of individual weather variables, and comparison with an independent SHAP-based attribution showed that the two approaches identify overlapping sets of influential variables while ordering them differently. Because this cluster-specific analysis is limited to the maize dataset and depends on weather-defined groups derived from aggregated environmental summaries, and because the functional annotation rests on positional proximity rather than experimental validation, these results are best interpreted as exploratory evidence of cluster-dependent ranking differences under the current pipeline. Overall, these findings support environment-associated marker ranking as a useful exploratory analysis direction, while future work can test the reproducibility of these patterns in larger datasets and under alternative environment-grouping strategies.

## References

[1] Ray, D. K., Gerber, J. S., MacDonald, G. K., and West, P. C. Climate variation explains a third of global crop yield variability. Nature Communications, 6(1):5989, 2015.

[2] Lobell, D. B., Schlenker, W., and Costa-Roberts, J. Climate trends and global crop production since 1980. Science, 333(6042):616–620, 2011. mmm

[3] Hickey, L. T., Hafeez, A. N., Robinson, H., Jackson, S. A., Leal-Bertioli, S. C. M., Tester, M., Gao, C., Godwin, I. D., Hayes, B. J., and Wulff, B. B. H. Breeding crops to feed 10 billion. Nature Biotechnology, 37(7):744–754, 2019.

[4] Meinshausen, N. and Bühlmann, P. Stability selection. Journal of the Royal Statistical Society: Series B (Statistical Methodology), 72(4):417–473, 2010.

[5] Crossa, J., Pérez-Rodríguez, P., Cuevas, J., Montesinos-López, O., Jarquín, D., De Los Campos, G., Burgueño, J., González-Camacho, J. M., Pérez-Elizalde, S., Beyene, Y., and Dreisigacker, S. Genomic selection in plant breeding: methods, models, and perspectives. Trends in Plant Science, 22(11):961–975, 2017.

[6] Heffner, E. L., Sorrells, M. E., and Jannink, J. L. Genomic selection for crop improvement. Crop Science, 49(1):1–12, 2009.

[7] Crossa, J., Campos, G. D., Pérez, P., Gianola, D., Burgueño, J., Araus, J. L., Makumbi, D., Singh, R. P., Dreisigacker, S., Yan, J., and Arief, V. Prediction of genetic values of quantitative traits in plant breeding using pedigree and molecular markers. Genetics, 186(2):713–724, 2010.

[8] Burgueño, J., de los Campos, G., Weigel, K., and Crossa, J. Genomic prediction of breeding values when modeling × genotype environment interaction using pedigree and dense molecular markers. Crop Science, 52(2):707–719, 2012.

[9] Jarquín, D., Crossa, J., Lacaze, X., Du Cheyron, P., Daucourt, J., Lorgeou, J., Piraux, F., Guerreiro, L., Pérez, P., Calus, M., and Burgueño, J. A reaction norm model for genomic selection using high-dimensional genomic and environmental data. Theoretical and Applied Genetics, 127(3):595–607, 2014.

[10] Heslot, N., Akdemir, D., Sorrells, M. E., and Jannink, J. L. Integrating environmental covariates and crop modeling into the genomic selection framework to predict genotype by environment interactions. Theoretical and Applied Genetics, 127(2):463–480, 2014.

[11] Malosetti, M., Ribaut, J. M., and van Eeuwijk, F. A. The statistical analysis of multienvironment data: modeling genotype-by-environment interaction and its genetic basis. Frontiers in Physiology, 4:44, 2013.

[12] McFarland, B. A., AlKhalifah, N., Bohn, M., Bubert, J., Buckler, E. S., Ciampitti, I., Edwards, J., Ertl, D., Gage, J. L., Falcon, C. M., and Flint-Garcia, S. Maize genomes to fields (G2F): 2014–2017 field seasons: genotype, phenotype, climatic, soil, and inbred ear image datasets. BMC Research Notes, 13(1):71, 2020.

[13] Jarquin, D., De Leon, N., Romay, C., Bohn, M., Buckler, E. S., Ciampitti, I., Edwards, J., Ertl, D., Flint-Garcia, S., Gore, M. A., and Graham, C. Utility of climatic information via combining ability models to improve genomic prediction for yield within the genomes to fields maize project. Frontiers in Genetics, 11:592769, 2021.

[14] Millet, E. J., Kruijer, W., Coupel-Ledru, A., Alvarez Prado, S., Cabrera-Bosquet, L., Lacube, S., Charcosset, A., Welcker, C., van Eeuwijk, F., and Tardieu, F. Genomic prediction of maize yield across European environmental conditions. Nature Genetics, 51(6):952–956, 2019.

[15] Abdellaoui, A., Yengo, L., Verweij, K. J. H., and Visscher, P. M. 15 years of GWAS discovery: realizing the promise. The American Journal of Human Genetics, 110(2):179–194, 2023.

[16] Montesinos-López, O. A., Crespo-Herrera, L., Pierre, C. S., Cano-Paez, B., Huerta-Prado,G. I., Mosqueda-González, B. A., Ramos-Pulido, S., Gerard, G., Alnowibet, K., Fritsche-Neto, R., and Montesinos-López, A. Feature engineering of environmental covariates improves plant genomic-enabled prediction. Frontiers in Plant Science, 15:1349569, 2024.

[17] Trevisan, B. A., Junqueira, V. S., Florêncio, B. D., Coelho, A. S., Marcatti, G. E., and Resende, R. T. A framework for building enviromics matrices in mixed models. arXiv preprint arXiv:2501.04147, 2025.

[18] Costa-Neto, G., Fritsche-Neto, R., and Crossa, J. Nonlinear kernels, dominance, and envirotyping data increase the accuracy of genome-based prediction in multi-environment trials. Heredity, 126(1):92–106, 2021.

[19] Gillberg, J., Marttinen, P., Mamitsuka, H., and Kaski, S. Modelling G× E with historical weather information improves genomic prediction in new environments. Bioinformatics, 35(20):4045–4052, 2019.

[20] Costa-Neto, G., Crespo-Herrera, L., Fradgley, N., Gardner, K., Bentley, A. R., Dreisigacker, S., Fritsche-Neto, R., Montesinos-López, O. A., and Crossa, J. Envirome-wide associations enhance multi-year genome-based prediction of historical wheat breeding data. G3, 13(2):jkac313, 2023.

[21] Hu, X., Carver, B. F., El-Kassaby, Y. A., Zhu, L., and Chen, C. Weighted kernels improve multi-environment genomic prediction. Heredity, 130(2):82–91, 2023.

[22] Brault, C., Conley, E. J., Read, A. C., Green, A. J., Glover, K. D., Cook, J. P., Gill, H. S., Fiedler, J. D., and Anderson, J. A. Improving genomic prediction for plant disease using environmental covariates. Plant Methods, 21(1):114, 2025.

[23] Gogna, A., Kamali, B., Wimmer, V., Schmidt, R. H., Rezaei, E. E., Eckhoff, W. M., Reif,J. C., and Zhao, Y. Predicting enviromically adapted varieties with big data. Genome Biology, 27(1):3, 2026.

[24] Onogi, A., Sekine, D., Kaga, A., Nakano, S., Yamada, T., Yu, J., and Ninomiya, S. A method for identifying environmental stimuli and genes responsible for genotype-by-environment interactions from a large-scale multi-environment data set. Frontiers in Genetics, 12:803636, 2021.

[25] Pham, H., Reisner, J., Swift, A., Olafsson, S., and Vardeman, S. Crop phenotype prediction using biclustering to explain genotype-by-environment interactions. Frontiers in Plant Science, 13:975976, 2022.

[26] Khaki, S. and Wang, L. Crop yield prediction using deep neural networks. Frontiers in Plant Science, 10:621, 2019.

[27] Fernandes, I. K., Vieira, C. C., Dias, K. O., and Fernandes, S. B. Using machine learning to combine genetic and environmental data for maize grain yield predictions across multienvironment trials. Theoretical and Applied Genetics, 137(8):189, 2024.

[28] Shook, J., Gangopadhyay, T., Wu, L., Ganapathysubramanian, B., Sarkar, S., and Singh,A. K. Crop yield prediction integrating genotype and weather variables using deep learning.PLOS ONE, 16(6):e0252402, 2021.

[29] Kick, D. R., Wallace, J. G., Schnable, J. C., Kolkman, J. M., Alaca, B., Beissinger, T. M., Edwards, J., Ertl, D., Flint-Garcia, S., Gage, J. L., and Hirsch, C. N. Yield prediction through integration of genetic, environment, and management data through deep learning. G3: Genes, Genomes, Genetics, 13(4):jkad006, 2023.

[30] Togninalli, M., Wang, X., Kucera, T., Shrestha, S., Juliana, P., Mondal, S., Pinto, F., Govindan, V., Crespo-Herrera, L., Huerta-Espino, J., and Singh, R. P. Multi-modal deep learning improves grain yield prediction in wheat breeding by fusing genomics and phenomics. Bioinformatics, 39(6):btad336, 2023.

[31] Jubair, S. and Domaratzki, M. Crop genomic selection with deep learning and environmental data: A survey. Frontiers in Artificial Intelligence, 5:1040295, 2023.

[32] Jubair, S., Tremblay-Savard, O., and Domaratzki, M. Gxenet: Novel fully connected neural network based approaches to incorporate G×E for predicting wheat yield. Artificial Intelligence in Agriculture, 8:60–76, 2023.

[33] Yao, Z., Yao, M., Wang, C., Li, K., Guo, J., Xiao, Y., Yan, J., and Liu, J. GEFormer: A genotype-environment interaction-based genomic prediction method that integrates the gating multilayer perceptron and linear attention mechanisms. Molecular Plant, 18(3):527–549, 2025.

[34] Wang, C., Zhang, D., Ma, Y., Zhao, Y., Liu, P., and Li, X. WheatGP, a genomic prediction method based on CNN and LSTM. Briefings in Bioinformatics, 26(2):bbaf191, 2025.

[35] Li, Y., Ren, S., Li, J., Lee, J., Wan, J., and Gan, X. MeNet: A mixed-effect deep neural network for multi-environment genomic prediction of agronomic traits. Plant Communications, 2026.

[36] Li, R., Zhang, D., Han, Y., Liu, Z., Zhang, Q., Zhang, Q., Wang, X., Pan, S., Sun, J., and Wang, K. Hybrid deep learning approaches for improved genomic prediction in crop breeding. Agriculture, 15(11):1171, 2025.

[37] Jubair, S., Tucker, J. R., Henderson, N., Hiebert, C. W., Badea, A., Domaratzki, M., and Fernando, W. D. GPTransformer: a transformer-based deep learning method for predicting Fusarium related traits in barley. Frontiers in Plant Science, 12:761402, 2021.

[38] Li, J., Luo, W., Yu, H., Huang, X., Ma, J., Li, S., Li, Y., and Gu, L. GViT-GP: injecting the genomic relationship matrix as an inductive bias into a vision transformer via cross-attention for genomic prediction. Frontiers in Genetics, 17:1758565, 2026.

[39] Wang, H., Yan, S., Wang, W., Chen, Y., Hong, J., He, Q., Diao, X., Lin, Y., Chen, Y., Cao, Y., and Guo, W. Cropformer: An interpretable deep learning framework for crop genomic prediction. Plant Communications, 6(3), 2025.

[40] Zhou, S., Cheng, K., Lv, L., Jiang, J., Zhou, S., Zhou, Y., Xu, Z., Huang, Q., Yang, H., Chen, L., and Xu, Y. CropARNet: A deep learning framework for crop genomic prediction with attention and residual modules. Crop Design, 100118, 2025.

[41] Rijal, K., Holmes, C. M., Petti, S., Reddy, G., Desai, M. M., and Mehta, P. Inferring genotype–phenotype maps using attention models. PNAS Nexus, 5(3):pgag046, 2026.

[42] Bai, L., Wang, K., Zhang, Q., Zhang, Q., Wang, X., Pan, S., Zhang, L., He, X., Li, R., Zhang, D., and Han, Y. A study of maize genotype–environment interaction based on deep K-means clustering neural network. Agriculture, 15(4):358, 2025.

[43] Saha, R., Morshedian, A., Sun, J., Duncan, R., and Domaratzki, M. Obscured-ensemble models for genomic prediction. PLOS ONE, 20(11):e0334239, 2025.

[44] He, X., Wang, K., Zhang, L., Zhang, D., Yang, F., Zhang, Q., Pan, S., Li, J., Bai, L., Sun, J., and Liu, Z. HGATGS: Hypergraph attention network for crop genomic selection. Agriculture, 15(4):409, 2025.

[45] Kihlman, R., Launonen, I., Sillanpää, M. J., and Waldmann, P. Sub-sampling graph neural networks for genomic prediction of quantitative phenotypes. G3: Genes, Genomes, Genetics, 14(11):jkae216, 2024.

[46] Wang, K., Han, Y., Zhang, Y., Zhang, Y., Wang, S., Yang, F., Liu, C., Zhang, D., Lu, T., Zhang, L., and Liu, Z. Maize yield prediction with trait-missing data via bipartite graph neural network. Frontiers in Plant Science, 15:1433552, 2024.

[47] Yang, F., Zhang, D., Zhang, Y., Zhang, Y., Han, Y., Zhang, Q., Zhang, Q., Zhang, C., Liu, Z., and Wang, K. Prediction of corn variety yield with attribute-missing data via graph neural network. Computers and Electronics in Agriculture, 211:108046, 2023.

[48] Gupta, A. and Singh, A. Agri-GNN: A novel genotypic-topological graph neural network framework built on GraphSAGE for optimized yield prediction. arXiv preprint arXiv:2310.13037, 2023.

[49] Morshedian, A. and Domaratzki, M. LSTM-attention-guided graph neural networks for integrated genotype–environment modeling in maize yield prediction. PLOS Computational Biology, 22(5):e1013729, 2026.

[50] Washburn, J. D., Varela, J. I., Xavier, A., Chen, Q., Ertl, D., Gage, J. L., Holland, J. B., Lima, D. C., Romay, M. C., Lopez-Cruz, M., and de los Campos, G. Global genotype by environment prediction competition reveals that diverse modeling strategies can deliver satisfactory maize yield estimates. Genetics, 229(2):iyae195, 2025.

[51] Sagae, V. S., Nascimento, M., Nascimento, A. C., Silva, F. L., and Jarquin, D. Impact of environmental covariates summarization on predictive ability in genomic selection. The Plant Genome, 19(1):e70194, 2026.

[52] NASA Langley Research Center. Prediction Of Worldwide Energy Resources (POWER) Project. Available at: https://power.larc.nasa.gov/, accessed 2026.

[53] Bush, W. S. and Moore, J. H. Chapter 11: Genome-wide association studies. PLOS Computational Biology, 8(12):e1002822, 2012.

[54] Huang, X. and Han, B. Natural variations and genome-wide association studies in crop plants. Annual Review of Plant Biology, 65:531–551, 2014.

[55] Zeng, T., Meng, Z., Yue, R., Lu, S., Li, W., Li, W., Meng, H., and Sun, Q. Genome-wide association analysis for yield-related traits in maize. BMC Plant Biology, 22(1):449, 2022.

[56] Tolley, S. A., Brito, L. F., Wang, D. R., and Tuinstra, M. R. Genomic prediction and association mapping of maize grain yield in multi-environment trials based on reaction norm models. Frontiers in Genetics, 14:1221751, 2023.

[57] Ma, J. and Cao, Y. Genetic dissection of grain yield of maize and yield-related traits through association mapping and genomic prediction. Frontiers in Plant Science, 12:690059, 2021.

[58] Diers, B. W., Specht, J., Rainey, K. M., Cregan, P., Song, Q., Ramasubramanian, V., Graef, G., Nelson, R., Schapaugh, W., Wang, D., and Shannon, G. Genetic architecture of soybean yield and agronomic traits. G3: Genes, Genomes, Genetics, 8(10):3367–3375, 2018.

[59] Nguyen, T. T., Huang, J. Z., Wu, Q., Nguyen, T. T., and Li, M. J. Genome-wide association data classification and SNPs selection using two-stage quality-based random forests. BMC Genomics, 16(Suppl. 2):S5, 2015.

[60] Szymczak, S., Holzinger, E., Dasgupta, A., Malley, J. D., Molloy, A. M., Mills, J. L., Brody, L. C., Stambolian, D., and Bailey-Wilson, J. E. r2VIM: A new variable selection method for random forests in genome-wide association studies. BioData Mining, 9(1):7, 2016.

[61] Ahmed, M., Seraj, R., and Islam, S. M. S. The k-means algorithm: A comprehensive survey and performance evaluation. Electronics, 9(8):1295, 2020.

[62] Rousseeuw, P. J. Silhouettes: a graphical aid to the interpretation and validation of cluster analysis. Journal of Computational and Applied Mathematics, 20:53–65, 1987.

[63] Lundberg, S. M. and Lee, S. I. A unified approach to interpreting model predictions. Advances in Neural Information Processing Systems, 30, 2017.

[64] Kendall, M. G. A new measure of rank correlation. Biometrika, 30(1–2):81–93, 1938.

[65] Hufford, M. B., Seetharam, A. S., Woodhouse, M. R., Chougule, K. M., Ou, S., Liu, J., Ricci, W. A., Guo, T., Olson, A., Qiu, Y., and Della Coletta, R. De novo assembly, annotation, and comparative analysis of 26 diverse maize genomes. Science, 373(6555):655–662, 2021.

[66] Aleksander SA, Balhoff J, Carbon S, Cherry JM, Drabkin HJ, Ebert D, Feuermann M, Gaudet P, Harris NL, Hill DP. The Gene Ontology knowledgebase in 2023. Genetics. 2023 May 2;224(1):iyad031.

[67] Paysan-Lafosse T, Blum M, Chuguransky S, Grego T, Pinto BL, Salazar GA, Bileschi ML, Bork P, Bridge A, Colwell L, Gough J. InterPro in 2022. Nucleic Acids Research. 2023 Jan 6;51(D1):D418–27.

[68] Mistry J, Chuguransky S, Williams L, Qureshi M, Salazar GA, Sonnhammer EL, Tosatto SC, Paladin L, Raj S, Richardson LJ, Finn RD. Pfam: The protein families database in 2021. Nucleic Acids Research. 2021 Jan 8;49(D1):D412–9.

[69] Naithani S, Gupta P, Preece J, D’Eustachio P, Elser JL, Garg P, Dikeman DA, Kiff J, Cook J, Olson A, Wei S. Plant Reactome: a knowledgebase and resource for comparative pathway analysis. Nucleic Acids Research. 2020 Jan 8;48(D1):D1093–103.

[70] Chen Y, Zein I, Brenner EA, Andersen JR, Landbeck M, Ouzunova M, Lübberstedt T. Polymorphisms in monolignol biosynthetic genes are associated with biomass yield and agronomic traits in European maize (Zea mays L.). BMC Plant Biology. 2010 Jan 15;10(1):12.

[71] McLaughlin JE, Tumer NE. Roles of non-specific lipid transfer proteins in plant defense: structural and functional perspectives. Frontiers in Fungal Biology. 2025 Sep 16;6:1640465.

[72] Liu, W. C., Song, R. F., Zheng, S. Q., Li, T. T., Zhang, B. L., Gao, X., and Lu, Y. T. Coordination of plant growth and abiotic stress responses by tryptophan synthase β subunit 1 through modulation of tryptophan and ABA homeostasis in Arabidopsis. Molecular Plant, 15(6):973–990, 2022.

[73] Lim CW, Yang SH, Shin KH, Lee SC, Kim SH. The AtLRK10L1.2, Arabidopsis ortholog of wheat LRK10, is involved in ABA-mediated signaling and drought resistance. Plant Cell Reports. 2015 Mar;34(3):447–55.

